# Symbioses of alvinocaridid shrimps from the South West Pacific: No chemosymbiotic diets but partially conserved gut microbiomes

**DOI:** 10.1101/2023.02.08.527621

**Authors:** Pierre Methou, Valérie Cueff-Gauchard, Loïc N. Michel, Nicolas Gayet, Florence Pradillon, Marie-Anne Cambon-Bonavita

## Abstract

*Rimicaris exoculata* shrimps from hydrothermal vent ecosystems are known to host dense epibiotic communities inside their enlarged heads and digestive systems. Conversely, other shrimps from the family, described as opportunistic feeders have received less attention. We examined the nutrition and bacterial communities colonizing “head” chambers and digestive systems of three other alvinocaridids – *Rimicaris variabilis*, *Nautilocaris saintlaurentae* and *Manuscaris* sp. – using a combination of electron microscopy, stable isotopes and sequencing approaches. Our observations inside “head” cavities and on mouthparts showed only a really low coverage of bacterial epibionts. In addition, no clear correlation between isotopic ratios and relative abundance of epibionts on mouthparts could be established among shrimp individuals. Altogether, these results suggest that none of these alvinocaridids rely on chemosynthetic epibionts as their main source of nutrition. Our analyses also revealed a substantial presence of several Firmicutes within the foreguts and midguts of these shrimps, which closest known lineages were systematically digestive epibionts associated with alvinocaridids, and more broadly from digestive systems of other crustaceans from marine and terrestrial ecosystems. Overall, our study opens new perspectives not only about chemosynthetic symbioses of vent shrimps, but more largely about digestive microbiomes with potential ancient and evolutionarily conserved bacterial partnerships among crustaceans.

## Introduction

Microbial symbioses are a ubiquitous phenomenon in nature, expanding physiological capabilities and ecological niches of organisms (McFall-Ngai *et al*., 2013). In many places, these associations constitute the structural base of ecosystems such as in hydrothermal vents. There, chemosynthetic symbioses with microorganisms using the chemical energy arising from vent fluid emissions are found in all invertebrates, establishing the foundation of lush faunal assemblages (Dubilier *et al*., 2008; Sogin and Leisch, 2020).

Among them, *Rimicaris exoculata* and *Rimicaris kairei* form large aggregations of thousands of individuals, gathered at the close vicinity of fluid emissions respectively in the Mid Atlantic Ridge and the Central Indian Ridge (Zbinden and Cambon-Bonavita, 2020). These shrimps host a complex community of cocci, rod-shaped and filamentous epibionts on the inner side of their enlarged cephalothorax, i.e. the branchiostegite, and on setae covering the surface of their hypertrophied mouthparts (Zbinden *et al*., 2004; Petersen *et al*., 2010; Methou, Hikosaka, *et al*., 2022). These communities comprise a wide diversity of chemosynthetic partners including *Campylobacterota*, *α-*, *γ-* and *ζ-Proteobacteria* as well as *Desulfobacterota* among others (Zbinden *et al*., 2008; Petersen *et al*., 2010; Guri *et al*., 2012; Jan *et al*., 2014; Jiang *et al*., 2020; Cambon-Bonavita *et al*., 2021; Methou, Hikosaka, *et al*., 2022), from which their hosts derive most of their nutrition (Polz *et al*., 1998; Gebruk *et al*., 2000; Van Dover, 2002; Methou *et al*., 2020) through direct transtegumental transfer of organic compounds (Ponsard *et al*., 2013). This diversity of bacterial partners reflects a diversity of metabolisms based on a wide range of energy sources (Jan *et al*., 2014; Jiang *et al*., 2020; Cambon-Bonavita *et al*., 2021) enabling these animals to thrive in vent fields with contrasting profiles of fluid chemistries.

Besides, *R. exoculata* and *R. kairei* shrimps harbour another community of resident epibionts within their digestive system (Zbinden and Cambon-Bonavita, 2003; Durand *et al*., 2010, 2015; Aubé *et al*., 2022; Guéganton *et al*., 2022; Qi *et al*., 2022). In *R. exoculata* this digestive symbiosis exhibits a clear partitioning between organs with several lineages of *Firmicutes* affiliated to *Mycoplasmatales* located in the foregut (oesophagus and stomach) and *Firmicutes* from the *Clostridia* class as well as *Candidatus* Rimicarispirillum which are long thin Deferribacterota inserted between microvilli in their midgut (Aubé *et al*., 2022; Guéganton *et al*., 2022). Unlike chemoautotrophic symbionts from the cephalothoracic cavity, these epibionts are heterotrophic and were hypothesized to complement their host diet and participate in its immunity (Aubé *et al*., 2022). To date, other bacterial lineages often found in the digestive microbiome of *R. exoculata* such as *Campylobacterota* and *Gammaproteobacteria* (Durand *et al*., 2010, 2015) were only observed as transient rod-shaped and cocci cells in its alimentary bolus (Guéganton *et al*., 2022).

These symbiotic communities of the cephalothoracic cavity and the foregut are renewed alongside their host exoskeleton at each moult, whereas those from the midgut are maintained throughout their adult life (Corbari *et al*., 2008; Guri *et al*., 2012). The constant renewal of their microhabitat coupled with an absence of similar or closely related lineages in the surrounding environment of their host, question the transmission pathways of the Mycoplasmatales located in the foregut (Durand *et al*., 2015). Similarly, the lack of geographic clustering of Deferribacterota epibionts in the midgut of *R. exoculata*, which are also absent from the environment, suggests a maternal inheritance (Durand *et al*., 2015). However, these lineages were never detected on their egg broods along the entire embryonic development (Guri *et al*., 2012; Methou *et al*., 2019).

Apart from *R. exoculata* and *R. kairei*, symbioses have been found in two other alvinocaridid species, *R. hybisae* from the Mid Cayman Rise and *R. chacei* from the Mid-Atlantic Ridge, which however display different trophic relations toward their symbiosis (Nye *et al*., 2012; Assié, 2016; Apremont *et al*., 2018). *R. chacei* shrimps lack an hypertrophied cephalothorax and are only partially dependent on their chemosynthetic symbiosis, with a mixed diet of symbiotrophy, bacterivory and scavenging (Gebruk *et al*., 2000; Methou *et al*., 2020). Their digestive system also hosts similar symbiotic communities than for *R. exoculata* with the same partitioning among foreguts and midguts (Apremont *et al*., 2018; Guéganton *et al*., 2022). On the other hand, *R. hybisae* shows more similarity with the ecology of *R. exoculata* and *R. kairei*, forming dense aggregates around chimneys and with an enlarged cephalothorax heavily colonized by epibionts (Nye *et al*., 2012; Streit *et al*., 2015). However recent evidences from gut contents and isotopic compositions of *R. hybisae* individuals distributed at the vent site periphery suggest they might have retained an ability to feed on other sources, including facultative carnivory (Versteegh *et al*., 2022), in addition to chemosynthetic bacterial sources (Streit *et al*., 2015).

In other alvinocaridids, nutritional strategies have been hypothesized to be mostly opportunistic and scavenging, with the use of several food sources including bacterial mats, detritus or the predation of small invertebrates (Gebruk *et al*., 2000; Stevens *et al*., 2008; Van Audenhaege *et al*., 2019; Suh *et al*., 2022). Based on stable isotope compositions, it was suggested that *Rimicaris variabilis* and *Manuscaris* sp. from the Manus Basin could either be conventional grazers/scavengers or feed on episymbiotic autotrophic bacteria in a similar fashion to *R. exoculata* (Van Audenhaege *et al*., 2019). Yet, no extensive study has investigated the microbial communities from their branchiostegites, mouthparts or digestive system so far.

Our study explores the bacterial communities colonizing cephalothoracic cavities and digestive systems of three alvinocaridid species from hydrothermal vents of South West Pacific basins – *Rimicaris variabilis*, *Nautilocaris saintlaurentae* and *Manuscaris* sp. –, as well as their nutrition, using a combination of electron microscopy, multiple stable isotopes and sequencing approaches. Our aim was to examine symbiotic relationships across alvinocaridid species with distinct ecologies to better understand the role and evolution of these symbioses. We address the following questions: 1) Do alvinocaridid species described as opportunistic feeders host epibiotic communities in their cephalothoracic cavity and/or their digestive system? 2) Do these potential epibiotic communities comprise similar or related bacterial lineages to epibionts of other *Rimicaris* species? 3) Can these alvinocaridid species rely, at least partially, on chemosynthetic symbionts for their nutrition?

## Materials and Methods

### Field sampling

Alvinocaridid shrimps were collected during the Futuna3 2012 and CHUBACARC 2019 oceanographic expeditions on board the R/V *L’Atalante* using a suction sampler manipulated by the HOV Nautile and the ROV Victor 6000 respectively. A total of 81 *Rimicaris variabilis* individuals were sampled from eight hydrothermal vent fields: Pacmanus and Susu Knolls in the Manus basin, La Scala in the Woodlark basin, Phoenix in the North Fiji basin, Fatu Kapa in the Futuna volcanic arc and Mangatolo, ABE and Tow Cam in the Lau basin (Figure 1). In addition, 25 *Nautilocaris saintlaurentae* individuals were also sampled at the Phoenix, Fatu Kapa and Tow Cam vent fields, as well as one *Manuscaris* sp. at the Pacmanus vent field. Specimens were identified morphologically and confirmed by genetic barcoding of their COI gene with specific primers for alvinocarids, using the protocol from (Methou *et al*., 2020). All sequences have been deposited in GenBank under accession numbers OQ363903 – OQ364004 (see Table S1 for sampling summary with associated individual ID). The 16S rRNA dataset is available in the NCBI SRA repository (submission identifier SUB12697284 and BioProject identifier PRJNA932596).

**Figure 1.**
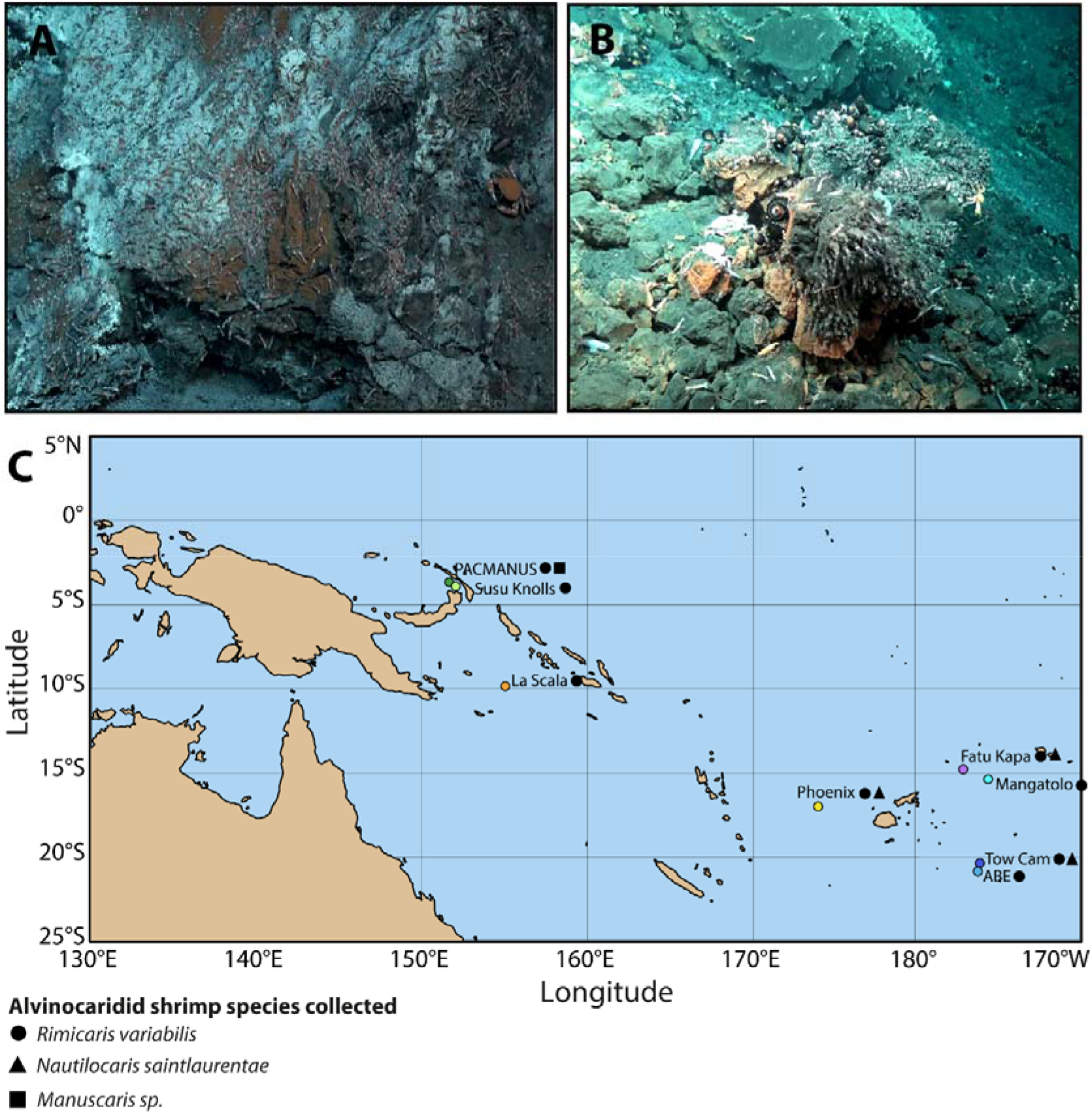
**A**. Alvinocaridids shrimps on the wall of an active vent chimney at Pacmanus (Manus basin) **B**. Alvinocaridids shrimps around assemblages of barnacles and at La Scala (Woodlark Basin) **C**. Sampling localities of alvinocaridid shrimps from Southwest Pacific basins. Colour dots depict hydrothermal vent field locations. Shapes depict shrimp species collected at a given sampling field.

Shrimps were dissected upon their recovery on board or in shore-based laboratory under sterile conditions to retrieve their anatomical parts: scaphognathites and exopodites mouthparts as well as branchiostegites from the cephalothoracic cavity, foreguts and midguts of their digestive system and pieces of abdominal muscles. Dissected parts or whole specimens were stored frozen at −80°C. Pieces from cephalothoracic cavities and mouthparts were also fixed in a 2.5% glutaraldehyde filtrated seawater solution for 16h at 4°C, rinsed and then stored at 4°C in filtrated seawater with 0,44 g/L of NaN_3_ at pH 7.4 until use for scanning electron microscopy (SEM) observations.

### Scanning Electron Microscopy

Dissected mouthparts and branchiostegites were dehydrated with an ethanol series (25, 50, 75, and 100% ethanol) and then for 5 h in a critical point dryer CPD 020 (Balzers Union, Balzers, Liechtenstein). Samples were then gold-coated with an SCD 040 (Balzers Union). Observations and imaging were performed using a Quanta 200 microscope (FEI-Thermo Fisher, Hillsboro, OR, United States).

### Stable isotope analysis

Abdominal muscle of alvinocaridid shrimps were oven-dried to constant mass at 50°C (>48 h) and ground into a homogeneous powder using a mortar and pestle. Measurements of stable isotope ratio were performed by continuous flow–elemental analysis–isotope ratio mass spectrometry (CF-EA-IRMS) at University of Liège (Belgium), using a vario MICRO cube C-N-S elemental analyser (Elementar Analysensysteme GMBH, Hanau, Germany) coupled to an IsoPrime100 isotope ratio mass spectrometer (Isoprime, Cheadle, United Kingdom). Isotopic ratios were expressed in ‰ using the widespread δ notation (Coplen, 2011) relative to the international references: Vienna Pee Dee Belemnite (for carbon), Atmospheric Air (for nitrogen) and Vienna Canyon Diablo Troilite (for sulfur). Primary analytical standards used for these analyses were the following: Sucrose (IAEA-C-6; δ^13^C=−10.8 ± 0.5‰; mean ± SD), ammonium sulfate (IAEA-N-2; δ^15^N= 20.4 ± 0.1‰; mean ± s.d.) and silver sulfide (IAEA-S-2; δ^34^S=22.6 ± 0.1‰; mean ± s.d.). A secondary analytical standard, Sulfanilic acid (Sigma-Aldrich; δ^13^C=−25.6 ± 0.4‰; δ^15^N=−0.13 ± 0.4‰; δ^34^S = 5.9 ± 0.5‰; means ± s.d.) was also used as well as an internal laboratory standard (seabass muscle). These standards were analysed interspersed among samples with one replicate of each standard every 15 analyses. Standard deviations on multi-batch replicate measurements of secondary and internal laboratory standards were 0.2‰ for δ^13^C and δ^15^N, and 0.4‰ for δ^34^S.

SIBER (Stable Isotope Bayesian Ellipses in R; Jackson et al. 2011) was used to explore ecological niches in an R 4.2.1 statistical environment (R Core Team, 2020). Two separate sets of standard ellipses were constructed: one with δ^13^C and δ^15^N data and another with δ^13^C and δ^34^S data. Areas of these ellipses were also estimated using Bayesian model (SEA_B_) with direct intergroup pairwise comparisons of SEA_B_. The model solutions were presented using credibility intervals of probability density function distributions. Areas of all ellipses were also estimated using the SEAc correction for small sample sizes, as outlined in (Jackson *et al*., 2011).

### DNA extraction and sequencing

Twenty-three *Rimicaris variabilis*, six *Nautilocaris saintlaurentae* and one *Manuscaris* sp. specimens were used for DNA extraction of their mouthparts, foreguts and midguts, as well as 17 additional *R. variabilis* and two additional *N. saintlaurentae* for mouthparts only, using the Nucleospin^®^ Soil Kit (Macherey-Nagel, Germany) following manufacturer’s instructions. Three blanks (i.e., a negative DNA extraction control) were also performed in parallel with DNA extractions of shrimp specimens.

For sequencing of the V3-V4 variable region of 16S rRNA (Fadrosh *et al*., 2014) using Illumina’s MiSeq technology, libraries were prepared using two successive PCR steps. (a) PCR1: samples were amplified in triplicate using the 341/785 primers (Herlemann *et al*., 2011) to generate a 450 bp fragment. Half of the P5 (CTTTCCCTACACGACGCTCTTCCGATCT) and P7 (GGAGTTCAGACGTGTGCTCTTCCGATCT) Illumina adapters were included to the 5⍰ part of the 341 forward and 785 reverse primers, respectively. PCR1 amplifications were performed in a final volume of 50 μl using 1 or 2 μL of DNA, 1.25 U of TaqCore polymerase (MP Biomedicals), standard Buffer with final 1.5 mM MgCl2, 0.5 mM of each dNTP and 0.2 μM of each primer under the following conditions: initial denaturation at 95⍰C for 5 min, followed by 35 cycles of 95⍰C for 30 s, 53⍰C for 30 s and 72⍰C for 1 min, and a final elongation step at 72⍰C for 6 min. (b) PCR2: the three PCR1 replicates of each sample were then pooled and sent to the GenoToul platform (GeT-BioPuce, INSA, Toulouse, France). Amplicons were first purified and dosed. Then they were used as templates for the PCR2 to which are added Illumina-tailed primers targeting the half of Illumina adapters P5 and P7 used in the first PCR and a unique index per sample. After purification, all amplicons were pooled in equimolar concentrations to be sequenced on an Illumina MiSeq system using paired-end sequencing with standard kit V3(250⍰bp⍰×⍰2).

### Metabarcoding analysis

A total of 16 817 628 raw reads across 109 samples, averaging 150 157 reads per sample, were analysed using the DADA2 pipeline (Callahan *et al*., 2016) in an R 4.2.1 statistical environment (R Core Team, 2020). Sequences were truncated to 250 bp for forward reads and to 240 bp for reverse reads based on the average quality scores. Additionally, reads displaying “N”, a quality score below 2, and/or more than 2 expected errors were discarded. The error model was trained using 1000 000 sequences before denoising, and chimeric sequences were removed based on a consensus approach before the paired ends were assembled. Contaminants were removed using blank controls with the MicroDecon R package (McKnight *et al*., 2019b).

The final data set contained 10 635 660 reads, with an average of 94 961 sequences per sample after quality filtering. Representative sequences were classified into taxonomic groups using the SILVA 138 database (Quast *et al*., 2013). Additional filtering on abundance was conducted at a threshold of 0.01% (Bokulich *et al*., 2013) to remove sequences containing non-biologically-relevant amplicon sequence variants (ASVs) (Breusing *et al*., 2022). ASVs affiliated with mitochondria sequences of alvinocaridid shrimps were also manually removed from the data set.

Visualization and statistical analyses of 16S rRNA bacterial diversity were performed using the Phyloseq (v. 1.4.0) (McMurdie and Holmes, 2013) and vegan (v. 2.6.2) (Oksanen *et al*., 2008) R packages. Alpha diversity across the 109 samples was explored with ASVs number for richness and Inverse Simpson Index for evenness. Differences in richness and evenness among categories (hosting organs, vent fields, alvinocaridid host species) were compared with Kruskal-Wallis tests followed by Dunn post-hoc tests. For Beta diversity, dataset was normalized to proportions (McKnight *et al*., 2019a) and analysed using Bray-Curtis distance matrices with the “distance” function (Phyloseq R package). Homogeneity between categories was tested with the “betadisper” function (vegan R package), and significant differences between categories were tested by permutational analysis of variance (PERMANOVA; 999 permutations) with the “adonis2” function (vegan R package). Constrained ordinations with stable isotopes ratios for each hosting organs were achieved by canonical analyses on the principal coordinates (CAP) using the “ordinate” (Phyloseq R package) and “scores” (vegan R package) functions.

## Results

### Scanning Electron Microscopy Observations

Observation of *Rimicaris variabilis* branchiostegites under Scanning Electron Microscopy (SEM) showed that the inner part of their cephalothoracic cavities were mostly devoid of bacterial colonization (Figure 2A, 2B). Conversely, a more abundant bacterial colonization was observed on the external surface of *R. variabilis* cephalothorax, in particular on setae aligned along their ventral side, which were covered by thick and thin filamentous bacteria (Figure 2C). In 7 out of the 12 *R. variabilis* individuals observed, single layered mats of rod-shaped bacteria were found on their cephalothorax inner surfaces, either on the anterior part facing mouthparts or on the posterior part facing the gills (Figure 2D). In some instances (2 out of 12 individuals), small spots of filamentous bacteria were localized on the most anterior part of the branchiostegites.

**Figure 2.**
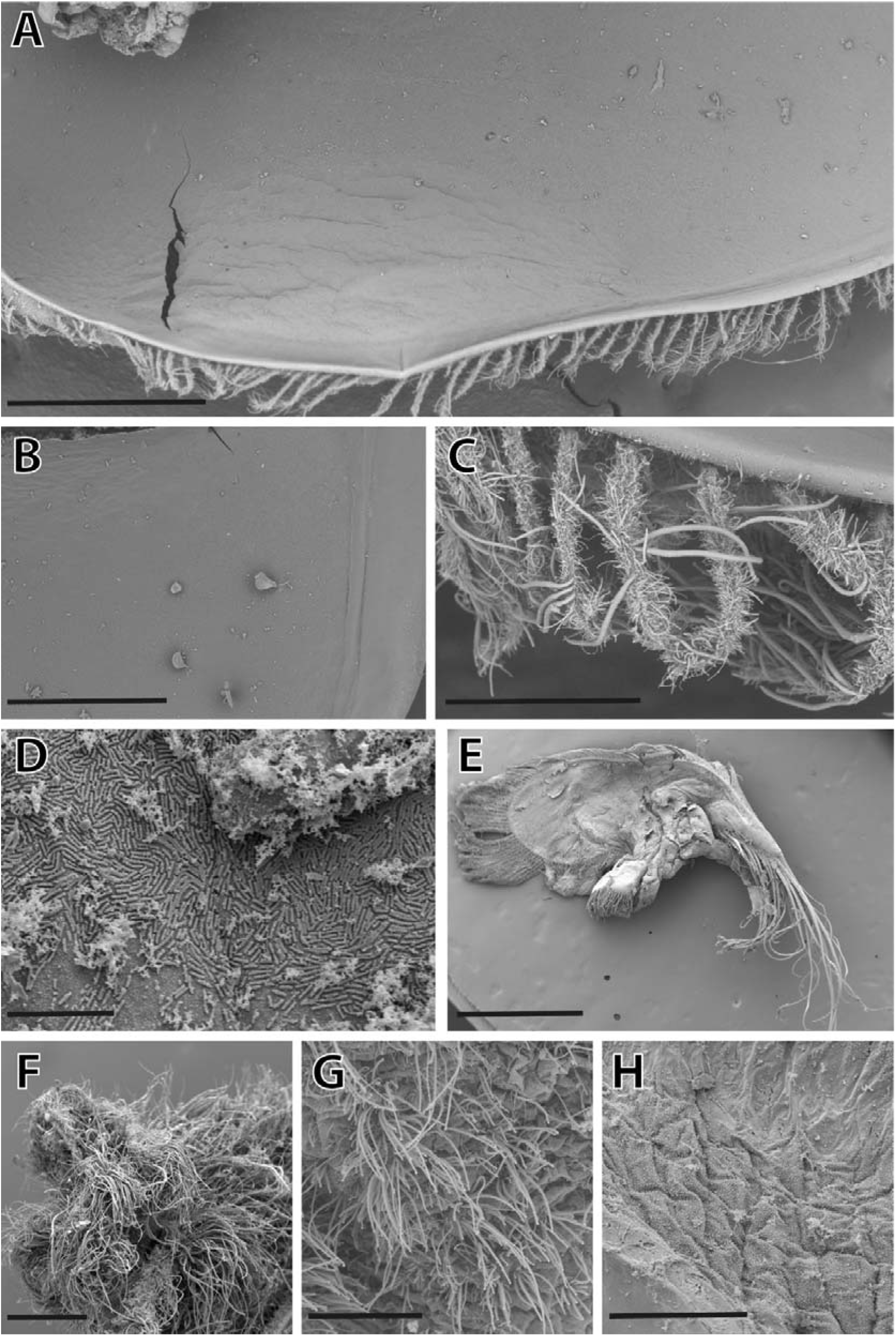
Scanning Electron Microscopy (SEM) observations of microbial communities on the surface of *Rimicaris variabilis* branchiostegites and mouthparts. **A**. Overview of the branchiostegite inner side. scale = 1 mm. **B**. Enlargement of the branchiostegite inner side devoid of bacterial colonization. scale = 500 μm. **C**. Filamentous bacteria colonizing the ventral setae along the external side of the branchiostegite. scale = 100 μm. **D**. Single-layered bacterial mats colonizing inner side of *R. variabilis* branchiostegite. scale = 10 μm. **E**. Overview of a scaphognathite dorsal side. scale = 1.5 mm. **F**. Dense aggregations of filamentous bacteria covering plumose setae of the scaphognathite margin. scale = 200 μm. **G**. Filamentous bacteria colonizing the scaphognathite surface. scale = 50 μm. **H**. Small cocci and rod-shaped bacteria colonizing the scaphognathite surface. scale = 50 μm.

Bacterial colonization was more widespread on *R. variabilis* mouthparts, although remaining limited to particular areas (Figure 2E). Dense aggregations of thick and thin filamentous bacteria covered plumose setae distributed along scaphognathite and exopodite margins (Figure 2F). Dorsal and ventral surfaces of these two mouthparts lacked bacteriophore setae (Figure 2E) and were generally only colonized by small cocci and rod-shaped bacteria on most of their surface (Figure 2G). In 2 out of 10 *R. variabilis* mouthparts observed, ventral and dorsal surfaces of scaphognathites were colonized by filamentous bacteria, but to a lesser extent than marginal setae (Figure 2H).

Observations of *Nautilocaris saintlaurentae* and *Manuscaris* sp. branchiostegites under SEM revealed similar patterns of bacterial colonization compared to *R. variabilis* with most of the inner parts of their cephalothoracic cavities devoid of bacteria (Figure 3A) or covered by single layered mats of rod-shaped bacteria (Figure 3B). As for *R. variabilis*, a few regionalized spots of filamentous bacteria were also present on the most anterior part of the *Manuscaris* sp. branchiostegite, close to the cephalothorax opening (Figure 3C). Bacterial colonization on mouthparts of these two species were also mostly limited to marginal setae covered by filamentous bacteria (Figure 3D), with only mono layers of cocci and rod-shaped bacteria on the surfaces of their scaphognathites and exopodites.

**Figure 3.**
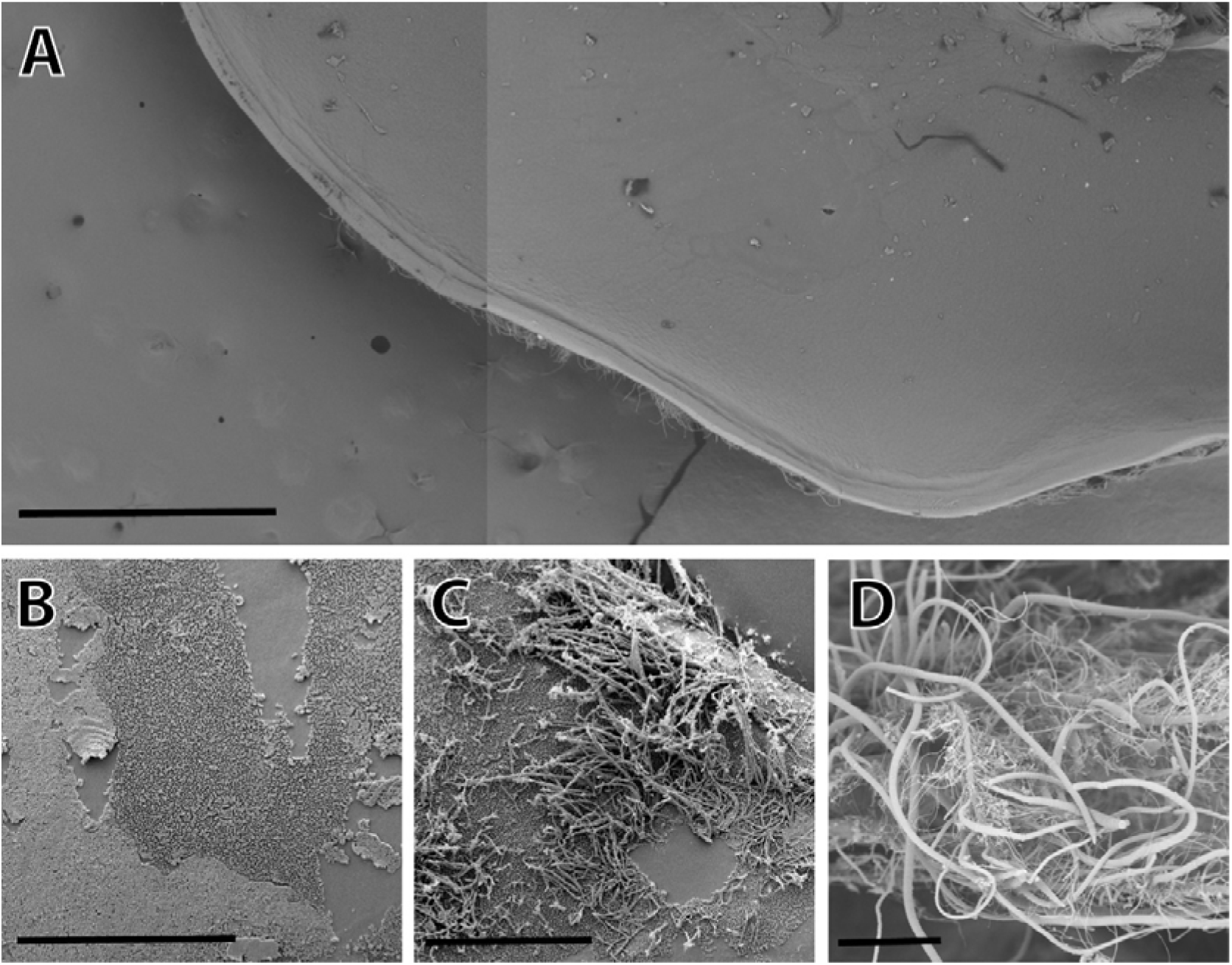
Scanning Electron Microscopy (SEM) observations of microbial communities on the surface of *Nautilocaris saintlaurentae* and *Manuscaris* sp. branchiostegites and mouthparts. **A**. Overview (composite image) of the inner side of *N. saintlaurentae* branchiostegite. scale = 1 mm. **B**. Single-layered bacterial mats colonizing inner side of *Manuscaris* sp. branchiostegite. scale = 50 μm. **C**. Spot of filamentous bacteria colonizing the most anterior part of the *Manuscaris* sp. branchiostegite. scale = 50 μm. **D**. Dense aggregations of filamentous bacteria covering plumose setae of *N. saintlaurentae* scaphognathite margin. scale = 50 μm.

### Stable Isotopes analysis

*Rimicaris variabilis* and *Nautilocaris saintlaurentae* populations showed limited variations in δ^13^C among vent fields (Kruskal–Wallis, p < 0.001; Figure S1A), with only significantly lower δ^13^C values for *R. variabilis* from Susu Knolls compared to those from Tow Cam, Pacmanus and La Scala (Dunn tests, p < 0.001; see Supplementary Table S2 for detailed p-values). Slight variations in δ^15^N among vent fields could be observed as well (Kruskal–Wallis, p < 0.001; Figure S1B) with higher δ^15^N values in *R. variabilis* from Pacmanus and La Scala compared to those from ABE, Fatu Kapa and Susu Knolls (Dunn tests, p < 0.001). Significant differences in δ^34^S of *Rimicaris variabilis* were also found among vent fields (Kruskal–Wallis, p < 0.001; Figure S1C) with a trend of ^34^S-depletion in shrimps from vent fields of the most eastern basins – Manus and Woodlark – compared to shrimp populations from more western basins – North Fiji and Lau – (Dunn tests, p < 0.001). At Fatu Kapa, δ^13^C, δ^15^N and δ^34^S of *R. variabilis* and *N. saintlaurentae* were similar between the two species (Dunn tests, p > 0.001).

SIBER analysis confirmed that carbon and sulfur isotopic niches of alvinocaridids from Manus and Woodlark basins were clearly separated from those of North Fiji and Lau populations (Figure 4A). However, the same trend was not observed for carbon and nitrogen isotopic niches (Figure 4B) with some overlap between *R. variabilis* from Pacmanus and Fatu Kapa (1.19‰^2^, i.e., 15.3% of the smallest ellipse area), from Tow Cam and Pacmanus (0.74‰^2^, i.e., 28.9% of the smallest ellipse area) or from Tow Cam and La Scala (0.31‰^2^, i.e., 12.1% of the smallest ellipse area). In general, limited overlap was observed between *R. variabilis* ellipses from vent fields within the same basin with no overlaps between the two Manus vent fields and small overlaps between vent fields from the Lau basin (22.4% of the smallest ellipse areas at most), except for carbon and sulfur ellipses of ABE and Fatu Kapa which were strongly overlapping (4.55‰^2^, i.e., 69.12% of the smallest ellipse area). Between Phoenix and Fatu Kapa, carbon and sulfur ellipses of *N. saintlaurentae* overlapped only by 0.64‰^2^, (i.e., 6.5% of the smallest ellipse area) but strongly overlapped for carbon and nitrogen ones (3.45 ‰^2^, i.e., 62.4% of the smallest ellipse area). At Fatu Kapa, ellipses of *R. variabilis* and *N. saintlaurentae* overlapped clearly, in particular for carbon and sulfur ellipses (6.7‰^2^, i.e., 57.7% of the smallest ellipse area) but also for carbon and nitrogen ellipses (2.02‰^2^, i.e., 36.6% of the smallest ellipse area).

**Figure 4.**
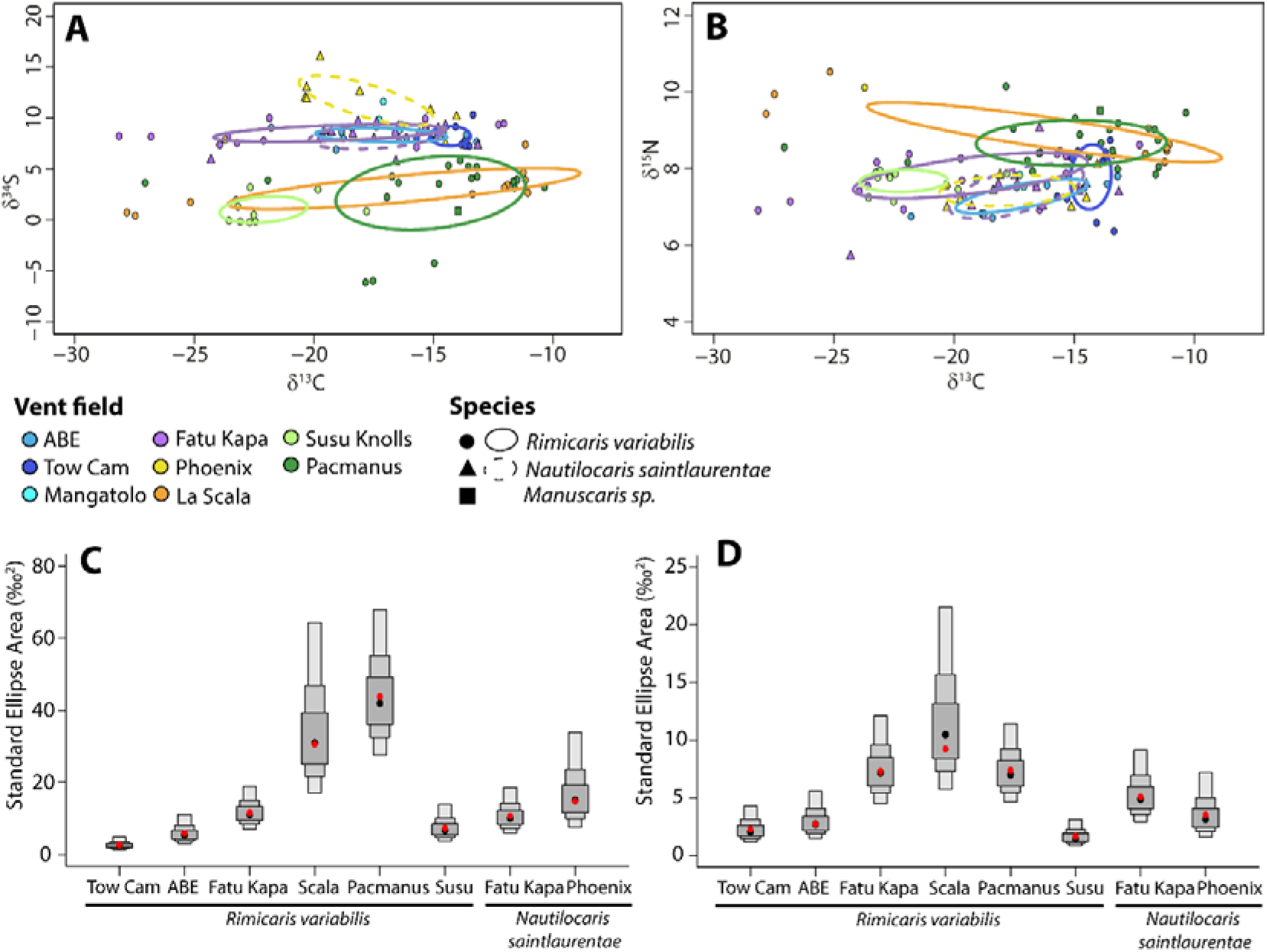
Isotopic niches of alvinocaridid shrimps from southwest Pacific basins. **A**. Carbon and sulfur isotopic niches **B**. Carbon and nitrogen isotopic niches. **A-B**. Each dot corresponds to the isotopic ratios of a shrimp individual; colors depict hydrothermal vent field locations and shapes depict different alvinocaridid species. **C**. Model-estimated bivariate standard area (SEA_B_) for carbon and sulfur ellipses **D**. Model-estimated bivariate standard area (SEA_B_) for carbon and sulfur ellipses **C-D**. Boxes in dark grey, medium grey, and light grey correspond, respectively, to the 50%, 75%, and 95% credibility intervals of probability density function distributions of the model solutions, and black dots are the modes of these distributions. Red dots are the standard ellipse areas computed using a frequentist algorithm adapted for small sample sizes (SEA_C_).

Areas of the standard ellipses associated with each shrimp species and vent field populations varied widely (Figure 4C and 4D), with SEA_c_ values ranging from 1.85‰^2^ (carbon and nitrogen ellipse of *R. variabilis* from Susu Knolls) to 46.24‰^2^ (carbon and sulfur ellipse of *R. variabilis* from Pacmanus). Overall, *R. variabilis* from Pacmanus had the widest isotopic niches (Figure 4C and 4D), with larger niches than any other shrimp populations in nearly all model solutions (>99.99% of model solutions for both carbon and sulfur niches and carbon and nitrogen niches) except for carbon and sulfur niches of *R. variabilis* from La Scala (only 72.09% of model solutions). These broad isotopic niches in some vent fields seemed to result in part from spatial variations, with in general, more similar and clustered isotopic ratios in individuals collected from the same sampling point, particularly at La Scala (Figure S2). Differences in niches sizes between alvinocaridid species at Fatu Kapa were not well supported by the model, with larger carbon and nitrogen niches for *R. variabilis* in 82.39% of model solutions and larger carbon and sulfur niches in 59.06% of model solutions.

### 16S rRNA metabarcoding analysis

Alpha Diversity analyses revealed slight variations in ASVs richness among host organs (Kruskal-Wallis, *H* = 11.65, *p* < 0.01) or among alvinocaridid species (Kruskal-Wallis, *H* = 7.06, *p* < 0.05) with a significantly higher number of ASVs in stomach compared to mouthparts communities (Dunn’s Multiple Comparison Test, *p* < 0.01) and a slightly higher number of ASVs in *Rimicaris variabilis* compared to *Nautilocaris saintlaurentae* communities (Dunn’s Multiple Comparison Test, *p* < 0.05). In contrast, ASVs richness was similar among back-arc basins (Kruskal-Wallis, *H* = 7.26, *p* > 0.05) or among vent fields (Kruskal-Wallis, *H* = 6.79, *p* > 0.05). Similarly, Inverse Simpson values did not indicate any variations of evenness among organs, shrimp species, regions or vent fields (Kruskal-Wallis tests, *p* > 0.05).

Based on PERMANOVA analyses, bacterial community composition was significantly influenced mostly by geography (i.e., among vent field; *F* = 4.832, *R*^2^ = 0.226, *p* < 0.001), but also by hosting organs (*F* = 6.005, *R*^2^ = 0.08, *p* < 0.001) or host species (*F* = 3.34, *R*^2^ = 0.045, *p* < 0.001). However, homogeneity of variances among vent fields (betadisper; *F* = 3.442, *p* < 0.01) or shrimp species (betadisper: *F* = 20.869, *p* < 0.001) were not met. Moreover, *R. variabilis* and *N. saintlaurentae* communities composition from Fatu Kapa and Phoenix taken alone – a balanced dataset (betadisper: *F* = 0.523, *p* > 0.05) – did not significantly differed between the two species (PERMANOVA: *F* = 1.752, *R*^2^ = 0.046, *p* > 0.05).

Canonical analysis on the principal coordinates (CAP) supported a correlation between stable isotopic composition of abdominal muscles and bacterial communities of alvinocaridid mouthparts (ANOVA-like: *F* = 3.042, *R*^2^ = 0.207, *p* < 0.001; Figure 5A) with a significant contribution of δ^15^N (*F* = 3.035, *p* < 0.01) and δ^34^S (*F* = 4.586, *p* < 0.001), but not δ^13^C (*F* = 1.504, *p* > 0.05). RDA models showed similar results for bacterial communities of alvinocaridid foreguts (ANOVA-like: *F* = 1.595, *R*^2^ = 0.184, *p* < 0.01; Figure 5A) and alvinocaridid midguts (ANOVA-like: *F* = 1.758, *R*^2^ = 0.203, *p* < 0.001; Figure 5C) with a significant contribution for δ^34^S but not or only slightly for δ^13^C on midguts and not for δ^15^N (see Supplementary Table 3). PERMANOVA analyses further confirmed that community compositions of each organ were mostly influenced by δ^34^S variations with only a significant influence of δ^15^N for bacterial communities of mouthparts and a slight effect of δ^13^C for bacterial communities of midguts (PERMANOVA tests; see Table S4 for detailed values).

**Figure 5.**
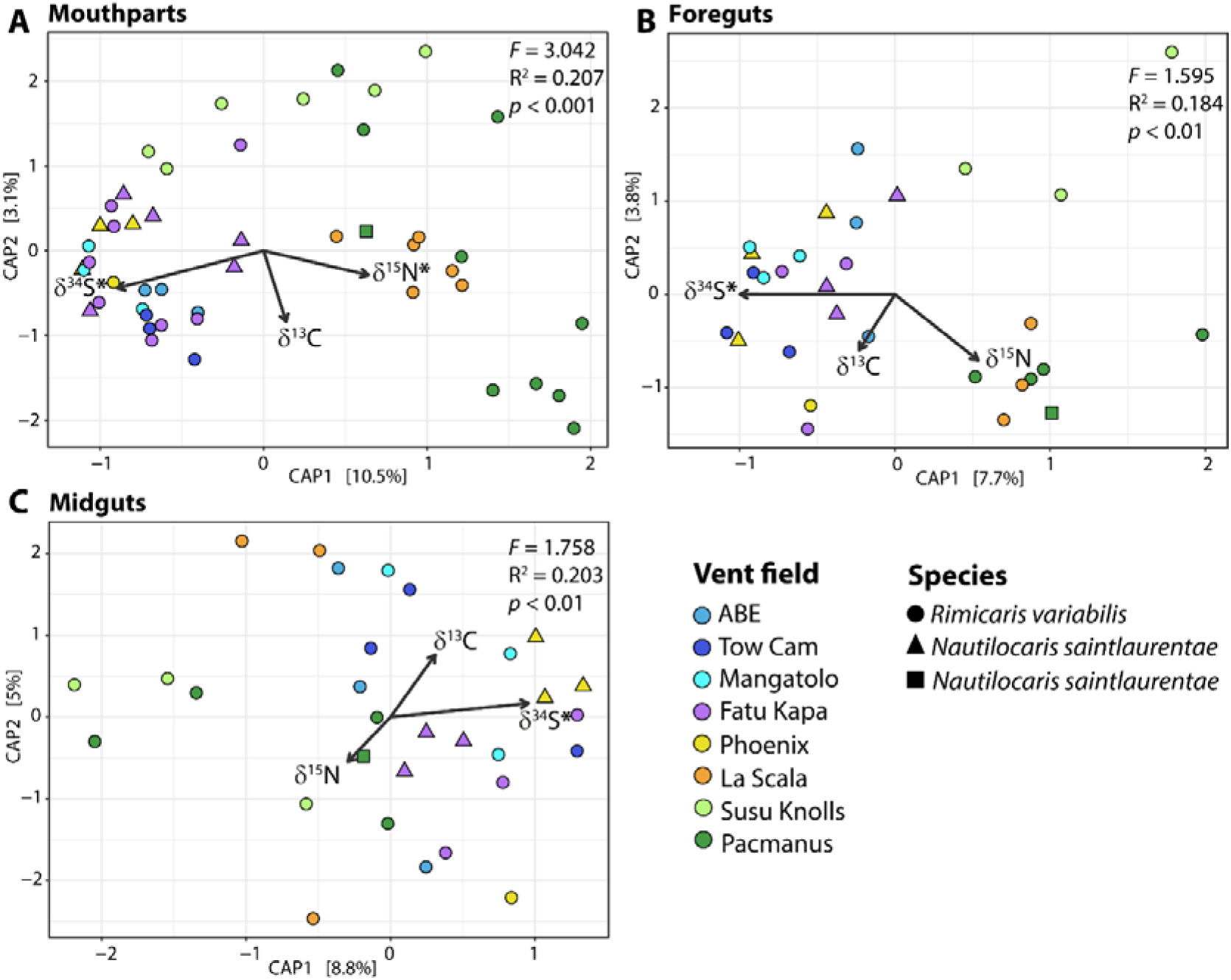
Constrained ordinations of 16S rRNA bacterial diversity by stable isotopes ratios using canonical analysis on the principal coordinates (CAP) for each hosting organs. **A**. Mouthparts bacterial communities **B**. Foreguts bacterial communities **C**. Midguts bacterial communities. Results from ANOVA-like permutation tests for CAP are displayed on each plot panel. Stable isotopic ratios which significantly contributed to CAP results are marked with an asterisk (p < 0.01, see supplementary Table 3). Points are coloured by hydrothermal vent field locations with shapes depicting distinct alvinocaridid species.

Composition of microbial communities on mouthparts were largely dominated by *Campylobacterota* ASVs (81.1% of mean relative abundance) followed by *Proteobacteria* ASVs (15.2%) (Figure 6A). They also included lower proportions of *Bacteroidota* ASVs (1.9%) and *Firmicutes* ASVs (1.1%); (Figure 6A). *Campylobacterota* ASVs also dominated microbial communities of foreguts (69.1%) and midguts (48.7%), but other groups had much higher relative abundances than on mouthparts, in particular *Firmicutes* (12.2% in foreguts and 31.1% in midguts respectively) but also *Verrucomicrobiota* (3.3% in foreguts and 2.8% in midguts respectively) and *Bacteroidota* to a lower extend (3.1% in foreguts and 6.1% in midguts respectively) (Figure 6B and 6C). In contrast, lower relative abundances of *Proteobacteria* were retrieved both in foreguts (8.6%) and in midguts (7.6%). Substantial proportions of *Desulfobacterota* were also found in foreguts (2.9%); (Figure 6B) and of *Fusobacteriota* in midguts (2.8%); (Figure 6C).

**Figure 6.**
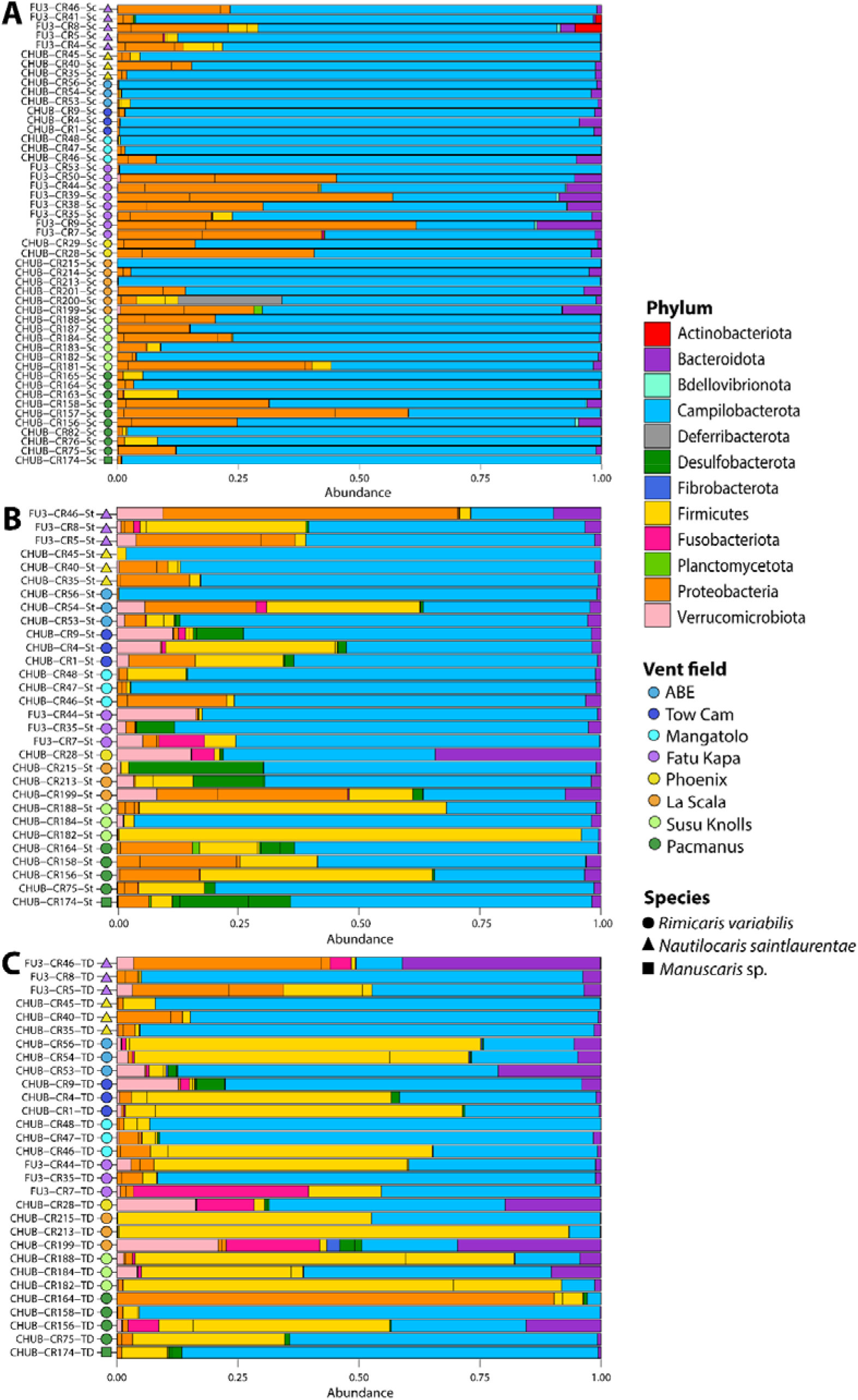
Relative abundances of 16S rRNA gene sequence reads from bacterial communities associated with southwest Pacific alvinocaridids according to their classification at the phylum level (Silva 138 database). **A**. Mouthparts bacterial communities **B**. Foreguts bacterial communities **C**. Midguts bacterial communities.

Phylogenetic reconstruction of *Firmicutes* ASVs agglomerated by phylogenetic similarity showed three main bacterial lineages in this phylum (Figure 7) with three ASVs affiliated to the *Candidatus* Hepatoplasmata genus (class Bacilli), three ASVs affiliated to the *Candidatus* Bacilloplasma genus (class Bacilli) and one ASV affiliated to the *Tyzzerella* genus (class *Clostridia*). Best BLAST hits of these ASVs were always *Rimicaris exoculata* or *Rimicaris chacei* foreguts and midguts epibionts with sequence similarity comprised between 98.7% and 99.9% (Figure 7 and Table S5). Most *Firmicutes* ASVs were present within each hosting organs and among each alvinocaridid species except ASV1723 that was found within *R. variabilis* foreguts and midguts only and ASV1649 that was only within *R. variabilis* and *N. saintlaurentae* foreguts and midguts (Figure 7).

**Figure 7.**
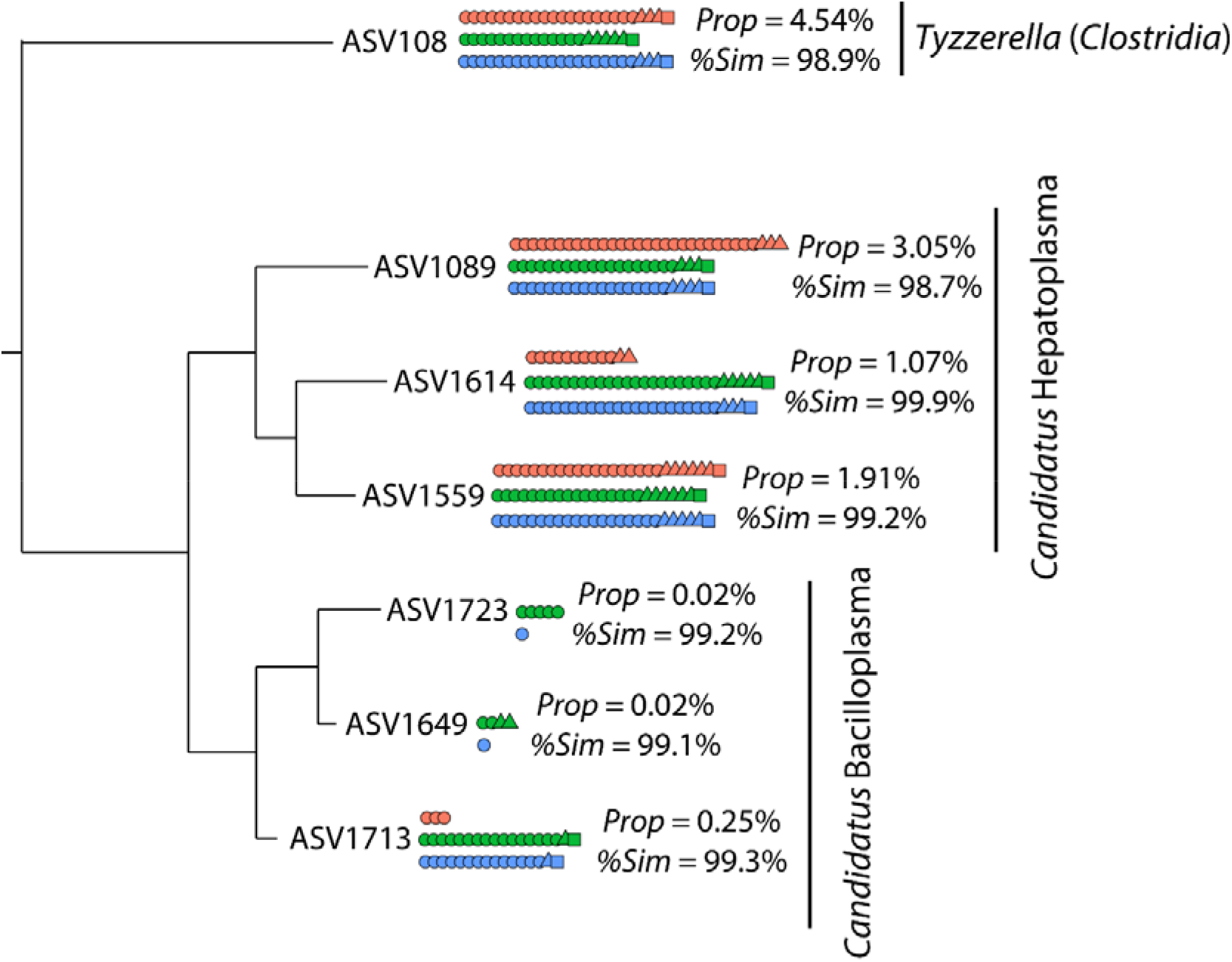
Phylogenetic tree of *Firmicutes* ASVs agglomerated by similarity (h = 0.1). The tree was constructed with the Maximum Likelihood method, based on the General Time Reversible model with Gamma distribution and allowing for some sites to be invariable (GTR+I+G). **Prop**: Relative abundance of the ASV among total sequence reads within the dataset. **%Sim**: % of similarity with the ASV best BLAST hit (*R. exoculata* or *R. chacei* digestive epibiont; details on Table S5). Each dot represent the occurrence of the lineage in an individual with shapes depicting alvinocaridid species (**circle**: *R. variabilis*; **triangle**: *N. saintlaurentae*; **square**: *Manuscaris* sp.) and colors depicting the hosting organ (**red**: mouthpart; **green**: stomach; **blue**: digestive tube).

## Discussion

### Nutritional strategies of alvinocaridid shrimps from hydrothermal vents of the Southwest Pacific basins

Our observations on the inner side of the cephalothoracic cavities showed only a scarce coverage of bacterial epibionts for either *Rimicaris variabilis*, *Nautilocaris saintlaurentae* or *Manuscaris* shrimps (Figure 2 & 3). A slightly more developed colonization was observed on their mouthparts with some filamentous bacteria although limited to particular areas, mostly the plumose setae on the mouthpart margins. This sharply contrasts with colonization patterns seen not only in alvinocaridid species relying mostly on their cephalothoracic chemosymbiosis such as *Rimicaris exoculata* (Zbinden *et al*., 2004; Petersen *et al*., 2010; Zbinden and Cambon-Bonavita, 2020) or *Rimicaris kairei* (Methou, Hikosaka, *et al*., 2022), but also in those with a mixed diet, only partially dependent on this symbiosis like *Rimicaris chacei* (Apremont *et al*., 2018), and which all exhibit extensive colonization of their cephalothoracic cavities by filamentous bacteria. Although earlier works stress out the importance of the moult cycle in *R. exoculata* symbiosis (Corbari *et al*., 2008), with a sparse colonization within the cephalothoracic cavity in early moult stage individuals, it is unlikely that moult stages have introduced a bias in our observations for alvinocaridids from South West pacific basins. Indeed, the inversely dense colonization of ventral setae along the external face of their cephalothorax (Figure 2C) suggests that limited colonization of their branchiostegites does not stem from a recent renewal of the exoskeleton but is found all along their moult cycle.

In addition, no clear correlation between the relative abundance of epibionts colonizing their mouthparts and stable isotopes ratios of carbon could be established among alvinocaridid individuals analysed in our study (Figure 5). In hydrothermal vent ecosystems, these variations in δ^13^C are mostly attributed to the use of different carbon fixation pathways by chemosynthetic microorganisms, with depleted δ^13^C ratios for those using the CBB cycle (typically −36 to −30‰) and enriched δ^13^C ratios for rTCA-fixed carbon sources (typically −15 to −10‰) (Hügler and Sievert, 2011; Portail *et al*., 2018). Both carbon fixation pathways can be found in chemosynthetic epibiont communities with *Campylobacterota* using the rTCA cycle and *Proteobacteria* using the CBB cycle (Jan *et al*., 2014; Jiang *et al*., 2020; Cambon-Bonavita *et al*., 2021). However, the relationship between individual δ^13^C ratios and relative abundance of rTCA- or CBB-fixing bacterial lineages did not hold for our dataset. As an example, the *R. variabilis* individual exhibiting the highest relative abundance of *Proteobacteria* lineages within its mouthparts (FU3-CR9; see Figure 5.) had a more enriched δ^13^C ratio (−16.6‰) compared to a *R. variabilis* individual from the same site (FU3-CR53; −23.6‰) whose mouthparts were completely dominated by *Campylobacterota* lineages and *Proteobacteria* being almost absent. Collectively, these results from microscopic observations, bacterial diversity and isotopic ratios all suggest that neither *R. variabilis*, *N. saintlaurentae* nor *Manuscaris sp*. rely on chemosynthetic epibionts as their main source of nutrition.

Aside a chemosymbiotic diet, other feeding modes such as bacterial grazers, scavengers or detritivores have been proposed for alvinocaridids shrimps, including those from the Manus and North Fiji basins (Van Audenhaege *et al*., 2019; Suh *et al*., 2022). Our results are consistent with previous studies on *R. variabilis* (Van Audenhaege *et al*., 2019; Suh *et al*., 2022), showing large trophic niches with particularly variable isotopic composition for carbon. Still, with a maximum of 11.6‰ for δ^34^S, their feeding sources remain within the range of chemosynthetically derived organic matter with no clear input of photosynthetic origin (Van Dover and Fry, 1994; Erickson *et al*., 2009; Reid *et al*., 2013). A notable exception were the *N. saintlaurentae* sampled at Phoenix site (North Fiji basin), with clearly higher δ^34^S ratio, up to 15.6‰, pointing out a potential mixed diet for this species with a larger contribution of photosynthetic material. However, the small size of these individuals and the observation of red lipid storages during their dissection (Methou, personal observation) rather indicate the influence of an ontogenetic shift as seen in juveniles and subadults stages of alvinocaridid shrimps from the Mid Atlantic Ridge (Pond *et al*., 1997; Methou *et al*., 2020), Central Indian Ridge (Van Dover, 2002) or the Mariana Arc (Stevens *et al*., 2008). Overall, these results suggest a generalist behaviour with various potential chemosynthetic food sources at the species level but more specialized feeding habits at a local scale. This is supported by the more similar and clustered isotopic ratios of shrimp individuals from the same sampling point within a vent field, arguing for a relatively strong habitat fidelity (Figure S2). Thus, although being potentially highly mobile animals, alvinocaridids from southwest Pacific might remain faithful to a same assemblage of foundation species once they settled, or at least at the timescales integrated by stable isotopic compositions of their abdominal muscles. Nevertheless, the presence of large alvinocaridid assemblages on chimney outcrops (Figure 1A), outside of mussel beds, tubeworm bushes, or hairy snail colonies, indicates that these shrimps do not solely rely on detritus of these foundational symbiotrophs, but are also able to feed on other nutrition sources, such as bacterial mats, possibly. Thereby, both detritivory and bacterivory diets could coexist in *R. variabilis* and *N. saintlaurentae*, although with strong intraspecific variations among individuals.

Interestingly, no isotopic niche partitioning was observed between *R. variabilis* and *N. saintlaurentae* from Fatu Kapa suggesting similar diets for the two co-occurring species (Figure 4). Since both were sometimes collected from the same assemblage, there was no clear indication of spatial segregation in distinct habitat either. To avoid competitive exclusion, niche theory predicts that sympatric species differ by their resource use and/or spatio-temporal habitat distribution, particularly in the case of closely related species with similar morphological traits and/or limited resource availability (Hutchinson, 1957; Schoener, 1974). However, the high biological productivity of hydrothermal vent ecosystems coupled with their temporal instability at short time scales might not allow to overreach the carrying capacity of these environments on the resources used by these shrimps, enabling long-term coexistence of similar species for the same food source. This contrasts with *Rimicaris* shrimps co-occurring in high densities assemblages on the Mid Atlantic Ridge, for which clear spatial and trophic niche partitioning could be observed between *R. exoculata* and *R. chacei* (Methou *et al*., 2020; Methou, Hernández-Ávila, *et al*., 2022). In the case of these species relying on their chemosymbiosis, competition for food is interlinked with competition for a limited space – i.e., the access to the vent fluid source – resulting ultimately in niche partitioning for the case of vent holobionts (Beinart *et al*., 2012; Van Audenhaege *et al*., 2019; Methou, Hernández-Ávila, *et al*., 2022). On the other hand, vent species with a distinct type of diet, such as alvinocaridids from the southwest Pacific might experience a more relaxed competition enabling co-occurring species to occupy the same ecological niche.

### Resident bacterial communities within the digestive system of alvinocaridid shrimps

Although bacterial coverage on mouthparts of *R. variabilis*, *N. saintlaurentae* and *Manuscaris* sp. was very low comparatively to *Rimicaris* species from the Atlantic or Indian Oceans, the composition of their epibiotic communities mirrors those previously observed in cephalothoracic cavities of the latter (Zbinden *et al*., 2008; Petersen *et al*., 2010; Guri *et al*., 2012; Jan *et al*., 2014; Apremont *et al*., 2018; Cambon-Bonavita *et al*., 2021; Methou, Hikosaka, *et al*., 2022). Thus, we found a similar phylogenetic diversity with a dominance of *Campylobacterota*, followed by several families of *Proteobacteria* - including *α-*, *γ-* and *ζ-proteobacteria* - as well as *Bacteroidota* epibionts.

In contrast, composition of their digestive communities differs, in part, from those of *R. exoculata* and *R. chacei* in the Mid Atlantic Ridge (Durand *et al*., 2010, 2015; Apremont *et al*., 2018; Aubé *et al*., 2022; Guéganton *et al*., 2022) or *R. kairei* in the Central Indian Ridge (Qi *et al*., 2022), which exhibited a clear partitioning of bacterial lineages between their digestive organs. Indeed, bacterial communities of their midguts were mainly composed of Deferribacterota and *Firmicutes* from the *Clostridia* class, whereas *Firmicutes* affiliated to *Mycoplasmatales* (class *Bacilli*) were dominant in foreguts (Durand *et al*., 2010; Aubé *et al*., 2022; Guéganton *et al*., 2022). These two phyla constitute resident communities within their respective hosting organs in *R. exoculata* and *R. chacei* whereas others bacterial lineages such as *Campylobacterota* or *Gammaproteobacteria* were only observed in the alimentary bolus (Guéganton *et al*., 2022). In the three species of alvinocaridids from the southwest Pacific, Deferribacterota were absent – except on the mouthpart of one *R. variabilis* individual from La Scala – and several lineages of *Firmicutes* affiliated to *Mycoplasmatales* and *Clostridia* were found in both the midguts and foreguts communities of every shrimp individual, often constituting the dominant lineage within their community (Figure 5B & 5C). These *Firmicutes* were also found on mouthparts of some individuals but in lower proportions than in their digestive systems (Figure 6A). This absence of partitioning among hosting organs in alvinocaridids from the Southwest Pacific is thus more similar to the case of the terrestrial isopod *Armadillidium vulgare* whose *Firmicutes* symbiont, *Candidatus* Hepatoplasma crinochetorum, occupying predominantly their hepatopancreas and caeca (Wang *et al*., 2004; Bouchon *et al*., 2016), was also found in other tissues, including their hindgut, nerve cord, gonads as well as their haemolymph and faeces (Dittmer *et al*., 2016).

It has been hypothesized that *Candidatus* Rimicarispirillum, the Deferribacterota symbionts of *R. exoculata*, supplement their host’s diet in vitamins through their biotin and riboflavin pathways, but depend on its supply for some essential amino acids (Aubé *et al*., 2022). The nature of the trophic diet in *R. variabilis*, *N. saintlaurentae* and *Manuscaris* sp. on the other hand, is quite different from that of *R. exoculata*, most likely detritivore and/or bacterivore (see discussion above), which could imply distinct needs to supplement their nutrition. Therefore, this apparent relationship between the presence of Deferribacterota symbionts in alvinocaridid microbiomes and their trophic strategies could suggest a tight nutritional link between the cephalothoracic and the digestive symbioses of these symbiotrophic animals.

Our results also reveal that the closest known lineages of *Firmicutes* found within the foreguts and midguts of southwest Pacific alvinocaridids were systematically epibionts associated with *R. exoculata* or *R. chacei* (Figure 7.) supporting a vertical inheritance of these symbionts and an association maintained along the evolutionary history of these hydrothermal vent shrimps. More broadly, related lineages like *Candidatus* Hepatoplasma or *Candidatus* Bacilloplasma were retrieved in digestive systems of several crustaceans such as terrestrial, intertidal or deep-sea isopods (Wang *et al*., 2004, 2016; Fraune and Zimmer, 2008; Eberl, 2010; Bouchon *et al*., 2016; Dittmer *et al*., 2016), hadal amphipods (Cheng *et al*., 2019), or coastal crab and shrimp species (Zhang *et al*., 2014, 2016; Chen *et al*., 2015). These *Candidatus* Hepatoplasma symbionts exhibited high level of specificity with their hosts in terrestrial isopods (Fraune and Zimmer, 2008). Taken together, these results would even suggest an ancient and evolutionarily conserved partnership in the crustacean subphylum. Conversely, the presence of each of these *Firmicutes* lineages within the digestive system of distantly-related alvinocaridids like *R. variabilis* and *N. saintlaurentae* but from the same geographic area (Figure 7.) is more congruent with a horizontal mode of transmission. Similarly, hepatopancreas of the co-occurring intertidal isopods, *Ligia pallasii* and *L. occidentalis*, hosted the same lineage of *Candidatus* Hepatoplasma (Eberl, 2010). It has been suggested that inter-moults and inter-generational transmission of *Mycoplasmatales* symbionts could be achieved by trophallaxis among individuals or by ingestion of their old cuticle (Durand *et al*., 2015). In the light of our results, this reinfection must be possible not only among individuals from the same species but also among individuals from the same family.

## Conclusion

Our study confirms that these opportunistic alvinocarids from the South West Pacific basins do not rely heavily on chemosymbiosis as an alternative or complementary part of their diet. Rather, they most likely feed on other food sources available at vent ecosystems, including bacterial mats, detritus or mucus discarded by the foundational symbiotroph species. On the other hand, part of their digestive microbiome, notably bacteria from the Firmicutes group, was highly conserved compared to other alvinocaridids but also more largely among crustaceans, suggesting overall a possible ancient and evolutionarily conserved bacterial partnership. However, distribution of these Firmicutes lineages within the different organs of the digestive system differs from those of other alvinocaridids where they are mostly restricted to the foregut. Of note, the almost absence of Deferribacterota residing in the digestive tube of alvinocaridids from other regions and exhibiting a different diet. A larger sampling comparing digestive microbiomes of different alvinocaridid species from several regions would be required to disentangle the respective influence of geography, host diet and host phylogeny of these associations.

## Acknowledgments

We thank the captain and crew of the French Research Vessel L’Atalante and to the team in charge of the ROV Victor 6000 during the ChubacArc expedition (2019) (https://doi.org/10.17600/18001111) and the HOV Nautile during the Futuna 3 expedition (2012) (https://doi.org/10.17600/12010040). Faunal collections were conducted with necessary authority permissions of the foreign countries. Permission for sampling in Exclusive Economic Zones (EEZ) was issued by the Papua New Guinea, The Republic of Fiji and Kingdom of Tonga. We obtained the agreement to sample in Wallis et Futuna waters by the Haut Commissariat à la République in New Caledonia and the Préfecture in Wallis and Futuna. Research animals were invertebrate caridean shrimps and no live experiments with animals were conducted in this study. We also are also grateful to the GenoToul platform (GeT-BioPuce, INSA, Toulouse, France) for providing resources for DNA sequencing.

## References

Apremont, V., Cambon-Bonavita, M.-A., Cueff-Gauchard, V., François, D., Pradillon, F., Corbari, L., and Zbinden, M. (2018) Gill chamber and gut microbial communities of the hydrothermal shrimp Rimicaris chacei Williams and Rona 1986: A possible symbiosis. PLoS One 13: e0206084.

Assié, A. (2016) Deep Se(a)quencing⍰: A study of deep sea ectosymbioses using next generation sequencing.

Aubé, J., Cambon-Bonavita, M.-A., Velo-Suárez, L., Cueff-Gauchard, V., Lesongeur, F., Guéganton, M., et al. (2022) A novel and dual digestive symbiosis scales up the nutrition and immune system of the holobiont Rimicaris exoculata. Microbiome 1–17.

Van Audenhaege, L., Fariñas-Bermejo, A., Schultz, T., and Lee Van Dover, C. (2019) An environmental baseline for food webs at deep-sea hydrothermal vents in Manus Basin (Papua New Guinea). Deep Res Part I Oceanogr Res Pap 148: 88–99.

Beinart, R.A., Sanders, J.G., Faure, B., Sylva, S.P., Lee, R.W., Becker, E.L., et al. (2012) Evidence for the role of endosymbionts in regional-scale habitat partitioning by hydrothermal vent symbioses. Proc Natl Acad Sci 241–250.

Bokulich, N.A., Subramanian, S., Faith, J.J., Gevers, D., Gordon, I., Knight, R., et al. (2013) Quality-filtering vastly improves diversity estimates from Illumina amplicon sequencing. Nat Methods 10: 57–59.

Bouchon, D., Zimmer, M., and Dittmer, J. (2016) The terrestrial isopod microbiome: An all-in-one toolbox for animal-microbe interactions of ecological relevance. Front Microbiol 7: 1–19.

Breusing, C., Castel, J., Yang, Y., Broquet, T., Sun, J., Jollivet, D., et al. (2022) Global 16S rRNA diversity of provannid snail endosymbionts from Indo-Pacific deep-sea hydrothermal vents. Environ Microbiol Rep.

Callahan, B.J., McMurdie, P.J., Rosen, M.J., Han, A.W., Johnson, A.J.A., and Holmes, S.P. (2016) DADA2: High-resolution sample inference from Illumina amplicon data. Nat Methods 13: 581–583.

Cambon-Bonavita, M.-A., Aubé, J., Cueff-Gauchard, V., and Reveillaud, J. (2021) Niche partitioning in the Rimicaris exoculata holobiont: the case of the first symbiotic Zetaproteobacteria. Microbiome 9: 1–16.

Chen, X., Di, P., Wang, H., Li, B., Pan, Y., Yan, S., and Wang, Y. (2015) Bacterial community associated with the intestinal tract of Chinese mitten crab (Eriocheir sinensis) farmed in Lake Tai, China. PLoS One 10: 1–21.

Cheng, X., Wang, Y., Li, J., Yan, G., and He, L. (2019) Comparative analysis of the gut microbial communities between two dominant amphipods from the Challenger Deep, Mariana Trench. Deep Res Part I Oceanogr Res Pap 103081.

Coplen, T.B. (2011) Guidelines and recommended terms for expression of stable-isotope-ratio and gas-ratio measurement results. Rapid Commun Mass Spectrom 25: 2538–2560.

Corbari, L., Zbinden, M., Cambon-Bonavita, M.A., Gaill, F., and Compère, P. (2008) Bacterial symbionts and mineral deposits in the branchial chamber of the hydrothermal vent shrimp rimicaris exoculata: Relationship to moult cycle. Aquat Biol 1: 225–238.

Dittmer, J., Lesobre, J., Moumen, B., and Bouchon, D. (2016) Host origin and tissue microhabitat shaping the microbiota of the terrestrial isopod Armadillidium vulgare. FEMS Microbiol Ecol 92: 1–15.

Van Dover, C.L. (2002) Trophic relationships among invertebrates at the Kairei hydrothermal vent field (Central Indian Ridge). Mar Biol 141: 761–772.

Van Dover, C.L. and Fry, B. (1994) Microorganisms as food resources at deep-sea hydrothermal vents. Limnol Oceanogr 39: 51–57.

Dubilier, N., Bergin, C., and Lott, C. (2008) Symbiotic diversity in marine animals: the art of harnessing chemosynthesis. Nat Rev Microbiol 6: 725–40.

Durand, L., Roumagnac, M., Cueff-Gauchard, V., Jan, C., Guri, M., Tessier, C., et al. (2015) Biogeographical distribution of Rimicaris exoculata resident gut epibiont communities along the Mid-Atlantic Ridge hydrothermal vent sites. FEMS Microbiol Ecol 91: 1–15.

Durand, L., Zbinden, M., Cueff-Gauchard, V., Duperron, S., Roussel, E.G., Shillito, B., and Cambon-Bonavita, M.A. (2010) Microbial diversity associated with the hydrothermal shrimp Rimicaris exoculata gut and occurrence of a resident microbial community. FEMS Microbiol Ecol 71: 291–303.

Eberl, R. (2010) Sea-land transitions in isopods: Pattern of symbiont distribution in two species of intertidal isopods Ligia pallasii and Ligia occidentalis in the Eastern Pacific. Symbiosis 51: 107–116.

Erickson, K.L., Macko, S.A., and Van Dover, C.L. (2009) Evidence for a chemoautotrophically based food web at inactive hydrothermal vents (Manus Basin). Deep Res Part II Top Stud Oceanogr 56: 1577–1585.

Fadrosh, D.W., Bing Ma, P.G., Sengamalay, N., Ott, S., Brotman, R.M., Ravel, J., et al. (2014) An improved dual-indexing approach for multiplexed 16S rRNA gene sequencing on the Illumina MiSeqplatform. Microbiome 2: 6.

Fraune, S. and Zimmer, M. (2008) Host-specificity of environmentally transmitted Mycoplasma-like isopod symbionts. Environ Microbiol 10: 2497–2504.

Gebruk, A. V., Southward, E.C., Kennedy, H., and Southward, A.J. (2000) Food sources, behaviour, and distribution of hydrothermal vent shrimps at the Mid-Atlantic Ridge. J Mar Biol Assoc United Kingdom 80: 485–499.

Guéganton, M., Rouxel, O., Durand, L., Cueff-Gauchard, V., Gayet, N., Pradillon, F., and Cambon-Bonavita, M.-A. (2022) Anatomy and Symbiosis of the digestive system of the vent shrimps Rimicaris exoculata and Rimicaris chacei revealed through imaging approaches. Front Mar Sci 7: 454–464.

Guri, M., Durand, L., Cueff-Gauchard, V., Zbinden, M., Crassous, P., Shillito, B., and Cambon-Bonavita, M.-A. (2012) Acquisition of epibiotic bacteria along the life cycle of the hydrothermal shrimp Rimicaris exoculata. ISME J 6: 597–609.

Herlemann, D.P.R., Labrenz, M., Jürgens, K., Bertilsson, S., Waniek, J.J., and Andersson, A.F. (2011) Transitions in bacterial communities along the 2000 km salinity gradient of the Baltic Sea. ISME J 5: 1571–1579.

Hügler, M. and Sievert, S.M. (2011) Beyond the Calvin Cycle⍰: Autotrophic Carbon Fixation in the Ocean. Ann Rev Mar Sci 3: 261–289.

Hutchinson, G. (1957) Concluding remarks. In Cold Spring Harbor Symposia on Quantitative Biology. pp. 66–77.

Jackson, A.L., Inger, R., Parnell, A.C., and Bearhop, S. (2011) Comparing isotopic niche widths among and within communities: SIBER - Stable Isotope Bayesian Ellipses in R. J Anim Ecol 80: 595–602.

Jan, C., Petersen, J.M., Werner, J., Teeling, H., Huang, S., Glöckner, F.O., et al. (2014) The gill chamber epibiosis of deep-sea shrimp Rimicaris exoculata: An in-depth metagenomic investigation and discovery of Zetaproteobacteria. Environ Microbiol 16: 2723–2738.

Jiang, L., Liu, X., Dong, C., Huang, Z., Cambon-Bonavita, M.-A., Alain, K., et al. (2020) “ Candidatus Desulfobulbus rimicarensis,” an Uncultivated Deltaproteobacterial Epibiont from the Deep-Sea Hydrothermal Vent Shrimp Rimicaris exoculata. Appl Environ Microbiol 86: 1–16.

McFall-Ngai, M., Hadfield, M.G., Bosch, T.C.G., Carey, H. V, Domazet-Lošo, T., Douglas, A.E., et al. (2013) Animals in a bacterial world, a new imperative for the life sciences. Proc Natl Acad Sci 110: 3229–3236.

McKnight, D.T., Huerlimann, R., Bower, D.S., Schwarzkopf, L., Alford, R.A., and Zenger, K.R. (2019a) Methods for normalizing microbiome data: An ecological perspective. Methods Ecol Evol 10: 389–400.

McKnight, D.T., Huerlimann, R., Bower, D.S., Schwarzkopf, L., Alford, R.A., and Zenger, K.R. (2019b) microDecon: A highly accurate read-subtraction tool for the post-sequencing removal of contamination in metabarcoding studies. Environ DNA 1: 14–25.

McMurdie, P.J. and Holmes, S. (2013) Phyloseq: An R Package for Reproducible Interactive Analysis and Graphics of Microbiome Census Data. PLoS One 8:.

Methou, P., Hernández-Ávila, I., Aube, J., Cueff-Gauchard, V., Gayet, N., Amand, L., et al. (2019) Is It First the Egg or the Shrimp? – Diversity and Variation in Microbial Communities Colonizing Broods of the Vent Shrimp Rimicaris exoculata During Embryonic Development. Front Microbiol 10: 1–19.

Methou, P., Hernández-Ávila, I., Cathalot, C., Cambon-Bonavita, M.-A., and Pradillon, F. (2022) Population structure and environmental niches of Rimicaris shrimps from the Mid-Atlantic Ridge. Mar Ecol Prog Ser 684: 1–20.

Methou, P., Hikosaka, M., Chen, C., Watanabe, H.K., Miyamoto, N., Makita, H., et al. (2022) Symbiont Community Composition in Rimicaris kairei Shrimps from Indian Ocean Vents with Notes on Mineralogy. Appl Environ Microbiol 88:.

Methou, P., Michel, L.N., Segonzac, M., Cambon-Bonavita, M.-A., and Pradillon, F. (2020) Integrative taxonomy revisits the ontogeny and trophic niches of Rimicaris vent shrimps. R Soc Open Sci 7: 200837.

Nye, V., Copley, J., and Plouviez, S. (2012) A new species of Rimicaris (Crustacea: Decapoda: Caridea: Alvinocarididae) from hydrothermal vent fields on the Mid-Cayman Spreading Centre, Caribbean. J Mar Biol Assoc United Kingdom 92: 1057–1072.

Oksanen, J., Kindt, R., Legendre, P., O’Hara, B., Simpson, G.L., Solymos, P.M., et al. (2008) The vegan package. Community Ecol Packag 190.

Petersen, J.M., Ramette, A., Lott, C., Cambon-Bonavita, M.-A., Zbinden, M., and Dubilier, N. (2010) Dual symbiosis of the vent shrimp Rimicaris exoculata with filamentous gamma- and epsilonproteobacteria at four Mid-Atlantic Ridge hydrothermal vent fields. Environ Microbiol 12: 2204–2218.

Polz, M.F., Robinson, J.J., Cavanaugh, C.M., Dover, C.L. Van, Van, C.L., and Van Dover, C.L. (1998) Trophic ecology of massive shrimp aggregations at a Mid-Atlantic Ridge hydrothermal vent site. Limnol Oceanogr 43: 1631–1638.

Pond, D.W., Segonzac, M., Bell, M. V., Dixon, D.R., Fallick, A.E., Sargent, J.R., et al. (1997) Lipid and lipid carbon stable isotope composition of the hydrothermal vent shrimp Mirocaris fortunata: Evidence for nutritional dependence on photosynthetically fixed carbon. Mar Ecol Prog Ser 157: 221–231.

Ponsard, J., Cambon-Bonavita, M.-A., Zbinden, M., Lepoint, G., Joassin, A., Corbari, L., et al. (2013) Inorganic carbon fixation by chemosynthetic ectosymbionts and nutritional transfers to the hydrothermal vent host-shrimp Rimicaris exoculata. ISME J 7: 96–109.

Portail, M., Brandily, C., Cathalot, C., Colaço, A., Gélinas, Y., Husson, B., et al. (2018) Food-web complexity across hydrothermal vents on the Azores triple junction. Deep Res Part I Oceanogr Res Pap 131: 101–120.

Qi, L., Lian, C.A., Zhu, F.C., Shi, M., and He, L.S. (2022) Comparative Analysis of Intestinal Microflora Between Two Developmental Stages of Rimicaris kairei, a Hydrothermal Shrimp From the Central Indian Ridge. Front Microbiol 12: 1–11.

Quast, C., Pruesse, E., Yilmaz, P., Gerken, J., Schweer, T., Yarza, P., et al. (2013) The SILVA ribosomal RNA gene database project: Improved data processing and web-based tools. Nucleic Acids Res 41: 590–596.

R Core Team,. (2020) R: A language and environment for statistical computing.

Reid, W.D.K., Sweeting, C.J., Wigham, B.D., Zwirglmaier, K., Hawkes, J.A., McGill, R.A.R., et al. (2013) Spatial Differences in East Scotia Ridge Hydrothermal Vent Food Webs: Influences of Chemistry, Microbiology and Predation on Trophodynamics. PLoS One 8: 1–11.

Schoener, T.W. (1974) Resouce partitioning in ecological communities: Research on how similar species divide resources helps. Science (80-) 185: 27–39.

Sogin, E.M. and Leisch, N. (2020) Chemosynthetic symbioses. Curr Biol 30: R1137–R1142.

Stevens, C.J., Limén, H., Pond, D.W., Gélinas, Y., and Juniper, S.K. (2008) Ontogenetic shifts in the trophic ecology of two alvinocaridid shrimp species at hydrothermal vents on the Mariana Arc, western Pacific Ocean. Mar Ecol Prog Ser 356: 225–237.

Streit, K., Bennett, S.A., Van Dover, C.L., and Coleman, M. (2015) Sources of organic carbon for Rimicaris hybisae: Tracing individual fatty acids at two hydrothermal vent fields in the Mid-Cayman rise. Deep Res Part I Oceanogr Res Pap 100: 13–20.

Suh, Y.J., Kim, M., Lee, W.-K., Yoon, H., Moon, I., Jung, J., and Ju, S.-J. (2022) Niche partitioning of hydrothermal vent fauna in the North Fiji Basin, Southwest Pacific inferred from stable isotopes. Mar Biol 1–15.

Versteegh, E.A.A., Dover, C.L. Van, Audenhaege, L. Van, and Coleman, M. (2022) Multiple nutritional strategies of hydrothermal vent shrimp (Rimicaris hybisae) assemblages at the Mid-Cayman Rise. Deep Res Part I.

Wang, Y., Huang, J.M., Wang, S.L., Gao, Z.M., Zhang, A.Q., Danchin, A., and He, L.S. (2016) Genomic characterization of symbiotic mycoplasmas from the stomach of deep-sea isopod bathynomus sp. Environ Microbiol 18: 2646–2659.

Wang, Y., Stingl, U., Anton-erxleben, F., Geisler, S., Brune, A., and Zimmer, M. (2004) “Candidatus Hepatoplasma crinochetorum,” a New, Stalk-Forming Lineage of Mollicutes Colonizing the Midgut Glands of a Terrestrial Isopod. Appl Environ Microbiol 70: 6166–6172.

Zbinden, M., Le Bris, N., Gaill, F., and Compère, P. (2004) Distribution of bacteria and associated minerals in the gill chamber of the vent shrimp Rimicaris exoculata and related biogeochemical processes. Mar Ecol Prog Ser 284: 237–251.

Zbinden, M. and Cambon-Bonavita, M. (2020) Biology and ecology of Rimicaris exoculata, a symbiotic shrimp from deep-sea hydrothermal vents. Mar Ecol Prog Ser 652: 187–222.

Zbinden, M. and Cambon-Bonavita, M.A. (2003) Occurrence of Deferribacterales and Entomoplasmatales in the deep-sea Alvinocarid shrimp Rimicaris exoculata gut. FEMS Microbiol Ecol 46: 23–30.

Zbinden, M., Shillito, B., Le Bris, N., de Villardi de Montlaur, C., Roussel, E., Guyot, F., et al. (2008) New insigths on the metabolic diversity among the epibiotic microbial communitiy of the hydrothermal shrimp Rimicaris exoculata. J Exp Mar Bio Ecol 359: 131–140.

Zhang, M., Sun, Y., Chen, K., Yu, N., Zhou, Z., Chen, L., et al. (2014) Characterization of the intestinal microbiota in Pacific white shrimp, Litopenaeus vannamei, fed diets with different lipid sources. Aquaculture 434: 449–455.

Zhang, M., Sun, Y., Chen, L., Cai, C., Qiao, F., Du, Z., and Li, E. (2016) Symbiotic bacteria in gills and guts of Chinese mitten crab (Eriocheir sinensis) differ from the free-living bacteria in water. PLoS One 11:.

