## supplemental_material.pdf for "Symbioses of alvinocaridid shrimps from the South West Pacific: No chemosymbiotic diets but partially conserved gut microbiomes"

### Supplementary material

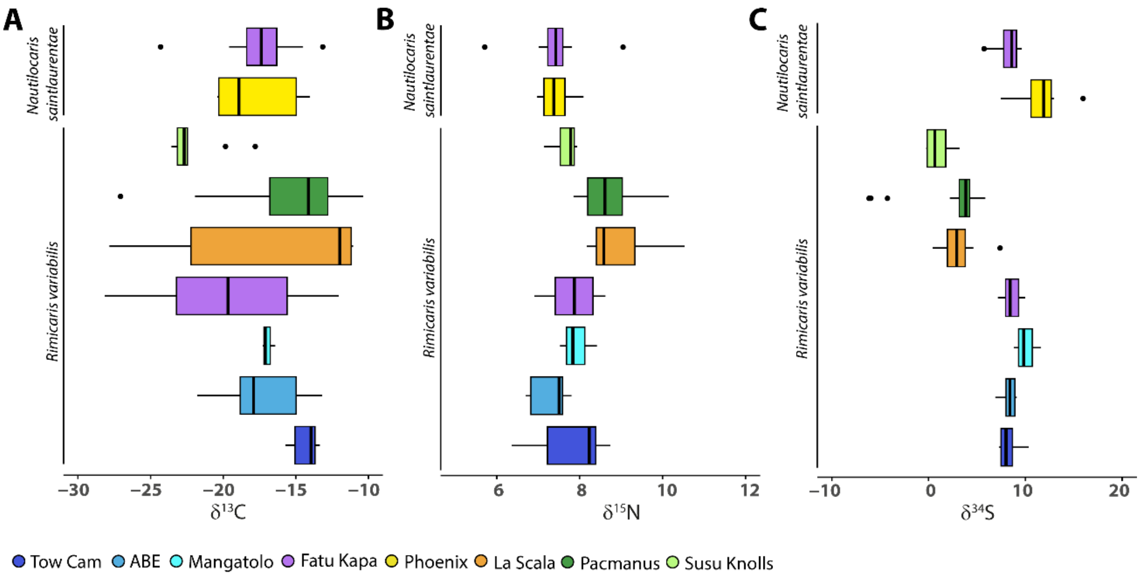


**Figure S1.** Isotopic ratios of alvinocaridids from Southwest Pacific basins for **A.** Carbon **B.** Nitrogen and **C.** sulfur.

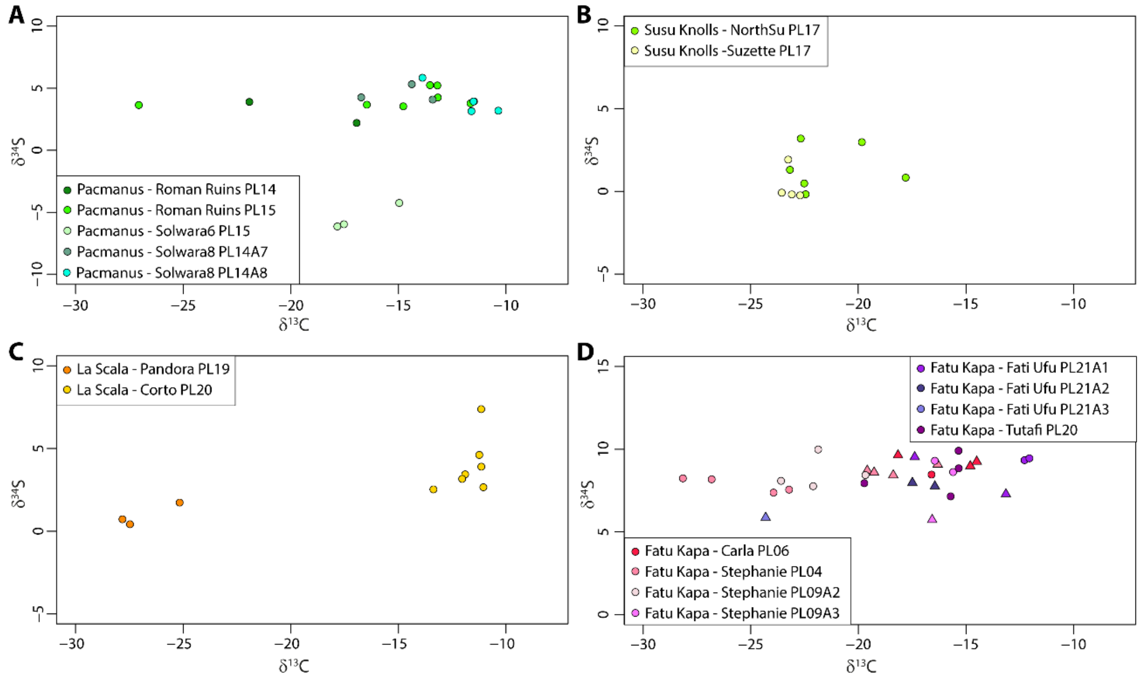


**Figure S2.** Carbon and sulfur isotopic ratios of alvinocaridid shrimps highlighting sampling events at **A**. the Pacmanus field **B.** Susu Knolls field **C.** La Scala field **D.** Fatu Kapa field.

**Table S1.** Summary of alvinocaridid shrimp sampling and conducted analyses with individual GenBank Identifier.

**Table S2.** p-values of Dunn tests used for isotopic ratio comparisons ($\delta^{13}C$, $\delta^{15}N$ $\delta^{34}S$) among alvinocaridid species and vent fields.

**Table S3.** Results from ANOVA-like permutation tests for RDA by term with 999 permutations for each hosting organs.

**Table S4.** Results from PERMANOVA tests of isotopic ratios with 999 permutations for each hosting organs.

**Table S5.** Best Blast hit results of Firmicutes ASVs from hosting organs of SW Pacific alvinocaridid shrimps.
